# Efficient and stable metabarcoding sequencing from DNBSEQ-G400 sequencer examined by large fungal community analysis

**DOI:** 10.1101/2020.07.02.185710

**Authors:** Xiaohuan Sun, Jingjing Wang, Chao Fang, Jiguang Li, Mo Han, Xiaofang Wei, Haotian Zheng, Xiaoqing Luo, Meihua Gong, Liang Xiao, Yuehua Hu, Zewei Song

**Affiliations:** BGI-Shenzhen, Shenzhen 518083, China; MGI, BGI-Shenzhen, Shenzhen 518083, China; CAS Key Laboratory of Tropical Forest Ecology, Xishuangbanna Tropical Botanical Garden, Chinese Academy of Sciences, Mengla 666303, China

**Author notes:** Correspondence should be addressed to Y.H. and Z.S.

## Abstract

Metabarcoding has become the de facto method for characterizing the structure of microbial communities in complex environmental samples. To determine how sequencing platform may influence microbial community characterization, we present a large-scale comparison of two sequencing platforms; Illumina MiSeq and a new platform DNBSEQ-G400 developed by MGI Tech. The accuracy of DNBSEQ-G400 on bacterial and fungal mock samples and compared sequencing consistency and precision between DNBSEQ-G400 and MiSeq platforms by sequencing the fungal ITS2 region from 1144 soil samples with 3 technical replicates. The DNBSEQ-G400 showed a high accuracy in reproducing mock communities containing different proportions of bacteria and fungi, respectively. The taxonomic profiles of the 1144 soil samples generated by the two DNBSEQ-G400 modes closely resembled each other and were highly correlated with those generated by the MiSeq platform. Analyses of technical replicates demonstrated a run bias against certain taxa on the MiSeq but not DNBSEQ-G400 platform. Based on lower cost, greater capacity, and less bias, we conclude that DNBSEQ-G400 is an optimal platform for short-term metabarcoding of microbial communities.

**IMPORTANCE:** Experimental steps that generate sequencing bias during amplicon sequencing have been intensively evaluated, including the choice of primer pair, polymerase, PCR cycle and technical replication. However, few studies have assessed the accuracy and precision of different sequencing platforms. Here, we compared the performance of newly released DNBSEQ-G400 sequencer with that of the commonly used Illumina MiSeq platform by leveraging amplicon sequencing of a large number of soil samples. Significant sequencing bias among major fungal genera was found in parallel MiSeq runs, which can be easily neglected without the use of sequencing controls. We emphasize the importance of technical controls in large-scale sequencing efforts and provide DNBSEQ-G400 as an alternative with increased sequencing capacity and more stable reproducibility for amplicon sequencing.

## INTRODUCTION

Culture independent studies have greatly expanded our knowledge of microbial diversity in recent decades (1–3). Studies focused on characterizing microbial community composition often amplify the 16S or ITS regions of rRNA (4, 5), which are the barcoding regions for bacteria and fungi, respectively. High throughput amplicon sequencing (HTS) allows for the parallel analysis of large numbers of samples from all kinds of natural environments, especially those with low microbial biomass, such as air (6), glacier (7), and deep-sea (8).

Factors contributing to ecological biases in HTS-based datasets have been reported in many studies, based on factors occurring both before (due to choice of polymerase, PCR primers and PCR cycles in library preparation (9, 10)) and after sequencing (due to choice of data analyzing methods (11–13)). In addition, using mock communities, run-to-run variation in taxon presence and abundance have also observed among 16S HTS using Illumina HiSeq (14). Similarly, bias among sequencing runs in detecting fungal taxa in soil samples has also reported (15), indicating the necessity of using positive controls in each sequencing event.

Compared to the intensive focus on biases in pre- and post-HTS processes, relatively few studies have focused on the potential bias introduced by sequencing platform. Currently, the dominant platforms used to generate HTS data of soil microbial communities is from Illumina (16, 17). Other sequencers, although wildly used in other areas, has not been recognized by researchers study microbial communities. In 2015, DNBSEQ™ platform was released by BGI using combinatorial Probe-Anchor Synthesis (cPAS) and DNA Nanoball (DNB) techniques (18). Since then DNBSEQ™ platform has been applied in various areas of genomics (18–20) and metagenomics (21, 22), with highly comparable quality to other platforms (23, 24). In 2017, an ultra-HTS sequencer DNBSEQ-G400, supporting multi-sequencing modes, including 2 × 200 paired-end (PE200) and 400 bases single end (SE400), and with a maximum throughput of 720 Gbp, was launched by MGI Tech. In comparison, Illumina’s MiSeq and NovaSeq generate 2 × 300 and 2 × 250 paired-end data, and with 15 Gbp and 400 Gbp maximum throughput respectively (Supplementary Table 1). With the comparable (and growing) read length and larger throughput, DNBSEQ-G400 appears to be a competitive candidate for microbial HTS analyses (Supplementary Table 1).

In this study, we examined the accuracy of DNBSEQ-G400 on bacterial and fungal mock samples and compared sequencing consistency and precision between DNBSEQ-G400 and MiSeq platforms by sequencing the fungal ITS2 region from 1144 soil samples with 3 technical replicates, respectively. The 1144 soil samples were sequenced in three DNBSEQ-G400 runs for both SE400 and PE200 modes, as well as eight runs using MiSeq. The 3 technical replicates, which were further assigned with 3 PCR replicates, i.e. 9 replicates, were sequenced in each run of the both platforms separately. Moreover, to assess performance consistency by a single sequencing mode within one platform, the 1144 samples were also repetitively sequenced two more times by SE400 mode only.

## RESULTS

### The accuracy amplicon sequencing on DNBSEQ platform

The performance of the DNBSEQ platform for HTS was initially examined by 16S V4 amplicons of the broadly used ZymoBIOMICS Microbial Community Standard and lab created combinations of multiple fungal species. The proportions of each taxon in the Zymo mock community were well reproduced (Fig. 1a), with no significant difference with the theoretical mock community relative abundance detected by pooled G-test of goodness-of-fit (p_value = 1). Four paired combinations of fungal taxa from ITS1 amplicons in three gradient proportions were also compared. Since genomic DNA usually contains an undefined number of rRNA copies, in order to quantify the accuracy between the amount of input and the results obtained by sequencing, we compared mixtures with the same proportions, using both raw genomic DNA and DNA following PCR amplification. Generally, DNA from PCR product mixtures reproduced the original proportions better than that from raw genomic DNA (Fig. 1b). Although no exact original proportion could be faithfully reproduced, taxon abundances showed similar gradients as those based on initial DNA ratios (Fig. 1b).

**Fig. 1.**
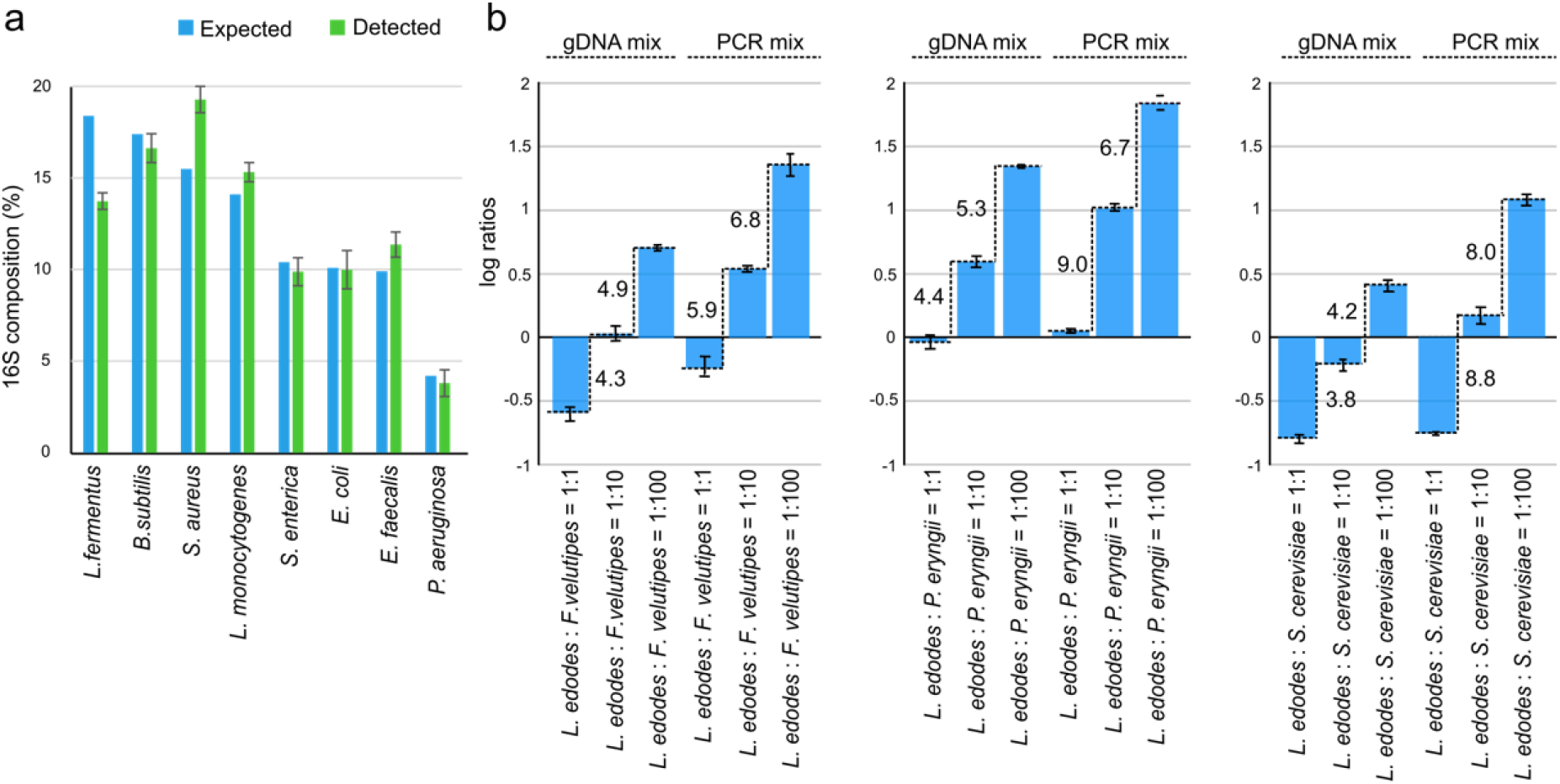
(A) Histogram of bacterial composition of ZymoBIOMICS Microbial Community Standards detected by DNBSEQ PE200 mode. (B) Histogram of log10 ratios of the proportions of mixtures of fungi sequenced by DNBSEQ SE400 mode. The numbers on the side of adjacent bars indicated the detected ratio of adjacent comparisons. The expected log ratios were shown with dotted lines. Error bars indicate 1 standard deviation.

### Consistency between DNBSEQ-G400 and MiSeq

Comparing ITS2 amplicons from 1144 soil samples between the DNBSEQ-G400 and MiSeq platforms revealed overall similarities regarding fungal community composition at the class level (Fig. 2, Supplementary Table 2). In total, 41 fungal classes were detected by MiSeq at the ALS forest plot, 38 and 37 had been identified by PE200 and SE400 modes of DNBSEQ-G400. At the LJ forest plot, 46, 42 and 43 fungal classes had been detected by MiSeq, PE200, and SE400, in which 39 were shared by the three modes. Finally, at the NBH forest plot, 37 overlapped classes were found among all the modes, and 37, 43 and 45 fungal genera had been identified by MiSeq, PE200 and SE400, respectively. Comparing to MiSeq PE300 and DNBSEQ-G400 SE400 modes, slightly fewer fungal genera were detected by DNBSEQ-G400 PE200 in all soil samples. Thirty-two fungal classes were identical across all platforms. Notably, the fungal classes that differed the most across platforms (11 out of 12 at ALS plot, 7 out of 8 at LJ, all 9 at NBH) had very low abundance (class abundance < 0.5%), with one only exception (class Archaeorhizomycetes, which was highly detected by PE200 at DNBSEQ platform).

**Fig. 2.**
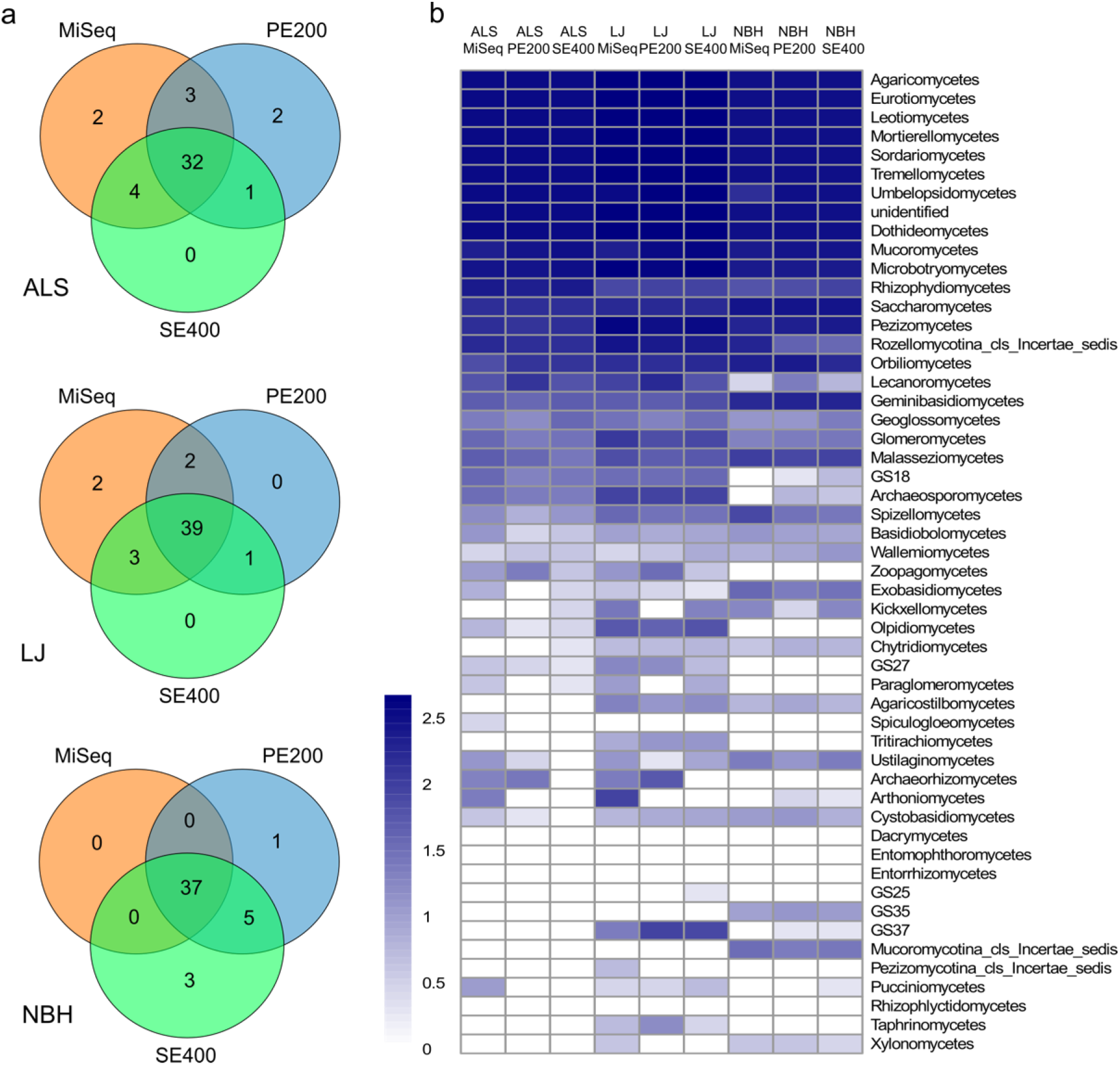
a) Venn diagrams show the overlap of identified fungal classes by MiSeq and DNBSEQ-G400 PE200 and SE400 modes. b) All classes from the three forest plots detected by MiSeq and DNBSEQ-G400 are represented as log_10_ (frequency) by heatmap. ALS=AiLaoShan, LJ=LiJiang, NBH=NaBanHe.

Similar to considerable overlap at the class taxonomic level, overall fungal community composition (based at the operational taxonomic unit (OTU) level) across the three different geographic locations were observed were not significant (p_value = 0.2223), examined by mixed linear model (Fig. 2b, supplementary Fig. 3). The frequency of occurrences of classes remained similar across the sequencing platforms showing by heatmap, especially for the top 20 most abundant classes (Fig. 2b). More variable classes, such as the Kickxellomycetes, Paraglomeromycetes, Archaeorhizomycetes and Arthoniomycetes, were again found at low abundant classes (Fig. 2b).

Sample-to-sample comparisons allowed us to further quantify the proportion difference of detected OTUs by the two sequencing platforms in a global scale. By using Procrustes analysis (25), we estimated the similarity of beta-diversity between two collections of samples, in our case, 1144 microbial community profiles from MiSeq and DNBSEQ-G400 (Fig. 3). The comparison of MiSeq to PE200 and SE400 both resulted in similar Procrustes profiles for all three plots, suggesting a closer correlation in the microbial taxa between PE200 and SE400. Indeed, higher correlation coefficient by permutation test was found within the two DNBSEQ-G400 sequencing modes (ALS=0.948, LJ=0.911, NBH=0.867) (Fig. 3). This high correlation was due to the high similarity of fungal communities between most PE200 and SE400 samples, with only a few samples showing high variance by Procrustes analysis. Moreover, the similarity between MiSeq and DNBSEQ-G400 differed associated with forest plots, which was higher for samples in LJ and NBH but lower for ALS (Fig. 3). We initially suspected that the performance difference might be related with species richness among the plots. However, ALS and LJ showed similar fungal diversities, as indicated by the median Shannon indexes of all soil samples (ALS = 2.85±0.44, LJ = 2.85±0.46), while NBH displayed the highest species diversity (NBH = 3.31±0.81). This indicates that species diversity was not the cause of the higher sequencing differences in ALS.

**Fig. 3.**
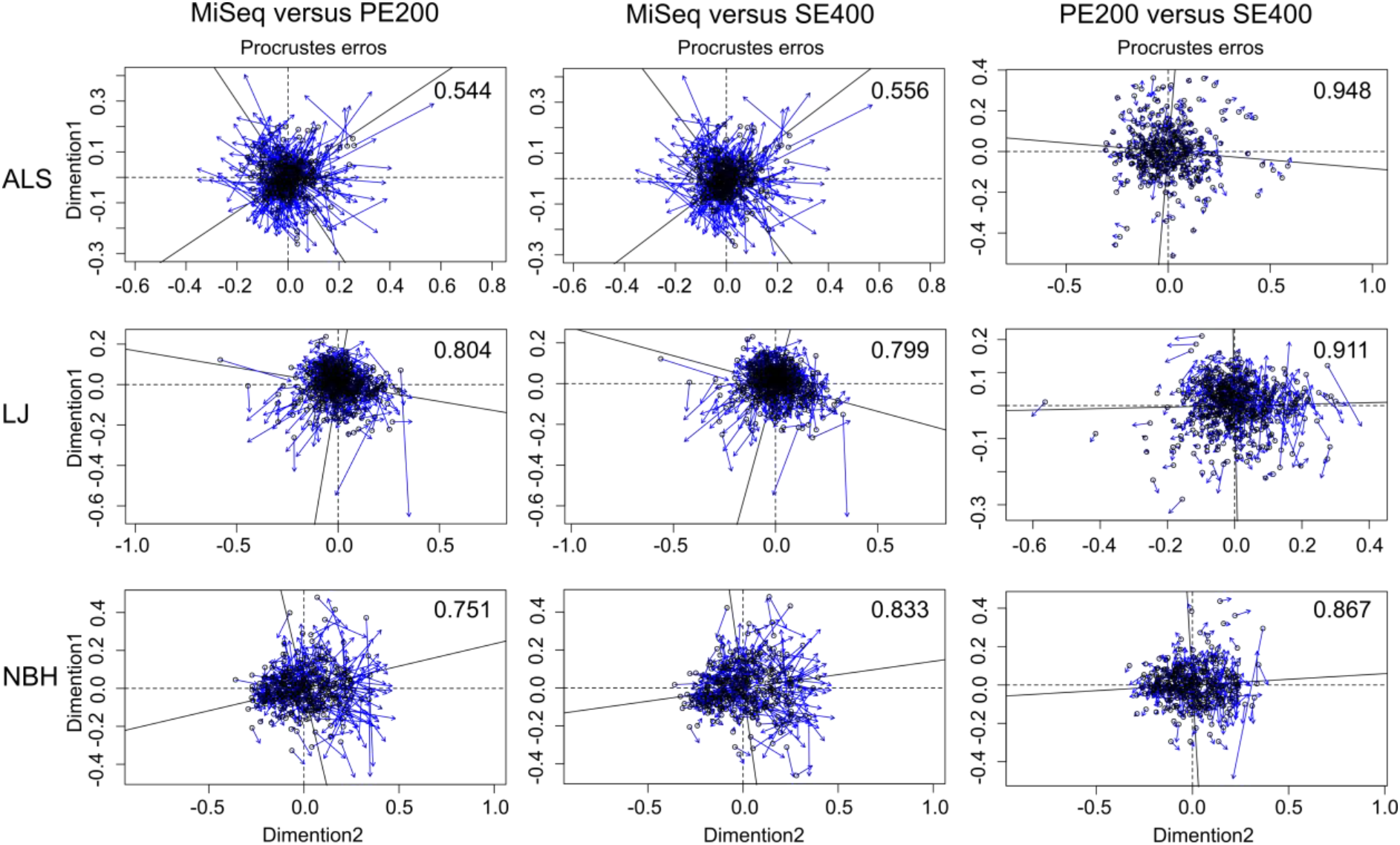
Procrustes analysis among Illumina MiSeq and DNBSEQ-G400 PE200 and SE400 sequencing platforms by sample-to-sample comparisons. Each sample contained a taxonomic set at species level. The round hollow circles showed the positions of multidimensional scaling conduced taxonomic profiles generated from one sequencing platform, and the blue arrows point to their positions generated by another sequencing platform. The correlation coefficients of platform comparisons are shown on the upper right corner of each figure. ALS=AiLaoShan, LJ=LiJiang, NBH=NaBanHe.

To further examine the sequencing consistency of DNBSEQ-G400, the same library of all 1144 soil samples had been further repetitively sequenced two more times by SE400 modes (SE400_1, SE400_2, SE400_3) with another DNBSEQ-G400 machine. A consistent performance of DNBSEQ-G400 was observed (Supplementary Fig. 4). The average correlation coefficients of fungal composition from the three sequencing repeats were 0.96, 0.99 and 0.99 for ALS, LJ and NBH, respectively. Further, the three repeats of all soil samples were clustered tightly by nonmetric Multidimensional Scaling (NMDS) analysis (Supplementary Fig. 4).

### Precision difference between DNBSEQ-G400 and MiSeq

To evaluate the differences in precision between DNBSEQ-G400 and MiSeq platforms, we performed NMDS ordination on the technical replicates, ALS268, LJ105 and NBH217 across all runs. On the MiSeq platform, technical PCR replicates were tightly clustered (Figure 4) but there was significant run-to-run variation, especially run4 and run6, indicating a bigger effect on precision caused by sequencing runs rather than PCR. These observations were supported by higher R^2^ value of samples organized by run (ALS268=0.85, LJ105=0.84, NBH217=0.75) compared to samples organized by PCR (ALS268=0.03, LJ105=0.03, NBH217=0.06) in nonparametric MANOVA analysis (Fig. 4a). In DNBSEQ-G400, run technical replicates from the same DNA sample were tightly clustered together (Fig. 4a), showing a consistent composition detection among DNBSEQ-G400 sequencing runs. Thus, the sequencing run exhibited a more extensive print on final fungal composition than that by PCR in MiSeq, while both PCR and run showed no significant effect in DNBSEQ-G400.

**Fig. 4.**
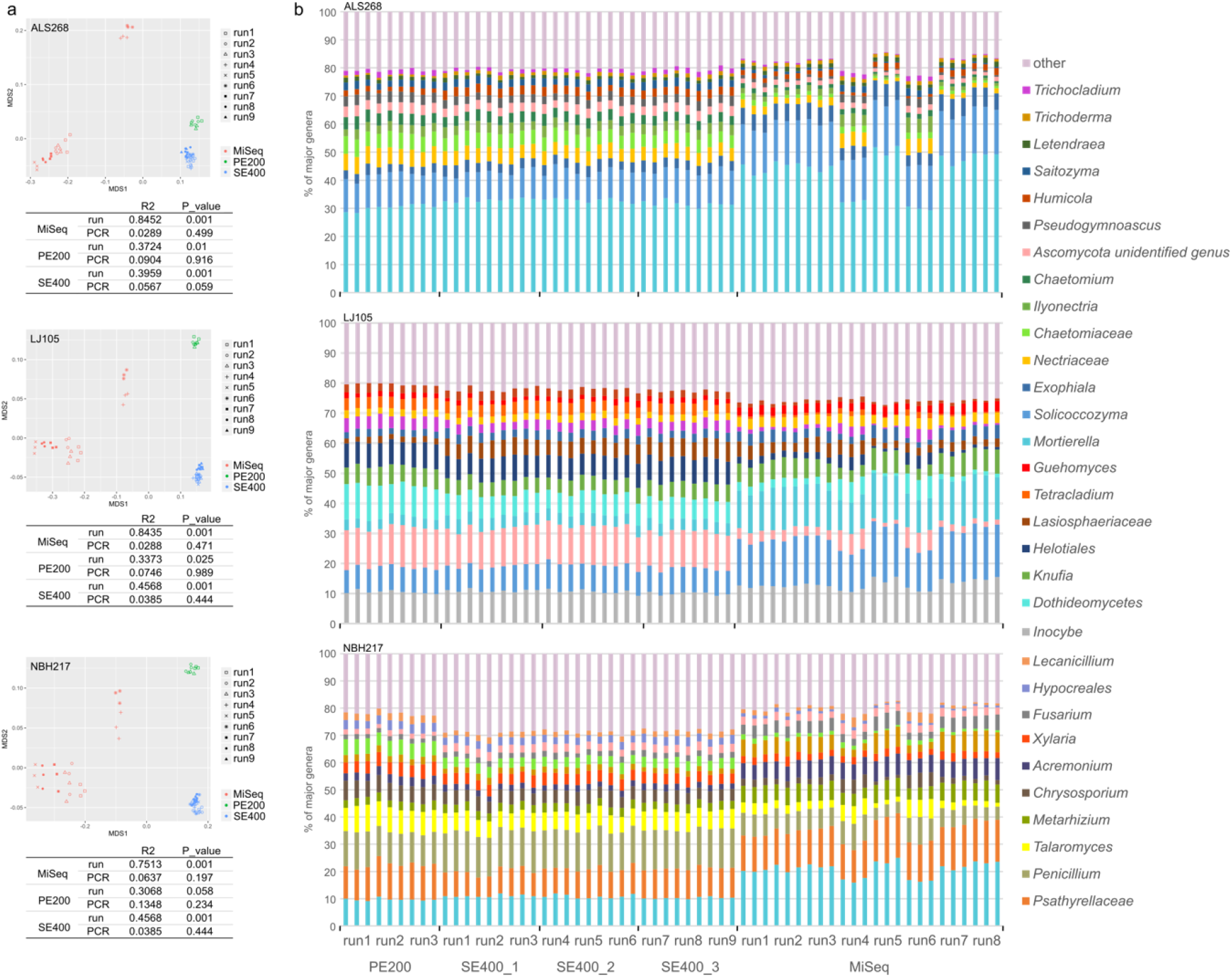
a) Non-metric multidimensional scaling (NMDS) analysis and ADONIS function illustrate the impact of sequencing run and PCR on the composition of fungal communities in three forest plots generated by DNBSEQ-G400 and MiSeq sequencing platforms. Samples indicate by labels with the same shape in NMDS are PCR replicates. b) bar chart shows the percentage of major OTUs in three forest plots at genus level in DNBSEQ-G400 and MiSeq sequencing platforms. ALS=AiLaoShan, LJ=LiJiang, NBH=NaBanHe.

The identities of major fungal genera of each technical replicate were very similar across all sequencing platforms but varied in proportion between DNBSEQ-G400 and MiSeq platforms (Fig. 4b). Nearly all major OTUs at Fig. 4b, except for the unidentified genus of Ascomycota from NBH and *Letendraea* and *Trichoderma* from ALS, exhibited significant differences in abundance (FDR < 0.001 at all three plots by Kruskal-Wallis test). Despite the differences among sequencers, within 8 MiSeq runs, run4 and run6 displayed different OTU abundance than other runs (Fig. 4b). Analysis of similarities (ANOSIM) showed significant differences within MiSeq 8 runs (p_value = 0.001 for technical replicates from all three plots). Spearman correlation coefficients between run4 or run6 and each other MiSeq runs were smaller than the correlation of pair-to-pair comparisons within other MiSeq runs. The relative weak correlation among run4 and run6 with other MiSeq runs was also reflected by larger p_value (0.01 < p_value < 0.05) than the pair comparisons among other runs (p_value < 0.01).

### OTUs with variable abundances detection

For those samples used as technical replicates, some OTUs exhibited strong variation among runs (Fig. 4b), indicating an important effect of sequencing run. OTUs with variable abundances were more frequently detected in the runs on the MiSeq platform (Fig. 5). An OTU showed significant abundance difference and low false discovery rate (FDR < 0.05) in at least one run compared with others was defined as a variable OTU (vOTUs). There were, accordingly, 50, 50 and 46 vOTUs in ALS268, LJ105 and NBH217, respectively. The vOTUs were always the highly abundant OTUs, as the vOTUs accounted for average 82% of the total fungal abundance in sample ALS268, 79% of total in LJ105, and 66% in NBH217 (Supplementary Table 3). vOTUs were enriched in Class *Sordariomycetes*, *Dothideomycetes*, *Eurotiomycetes*, *Tremellomycetes*, *Agaricomycetes* and *Mortierellomycetes*, which were also the principal fungal classes in the three forests (Fig. 5).

**Fig. 5.**
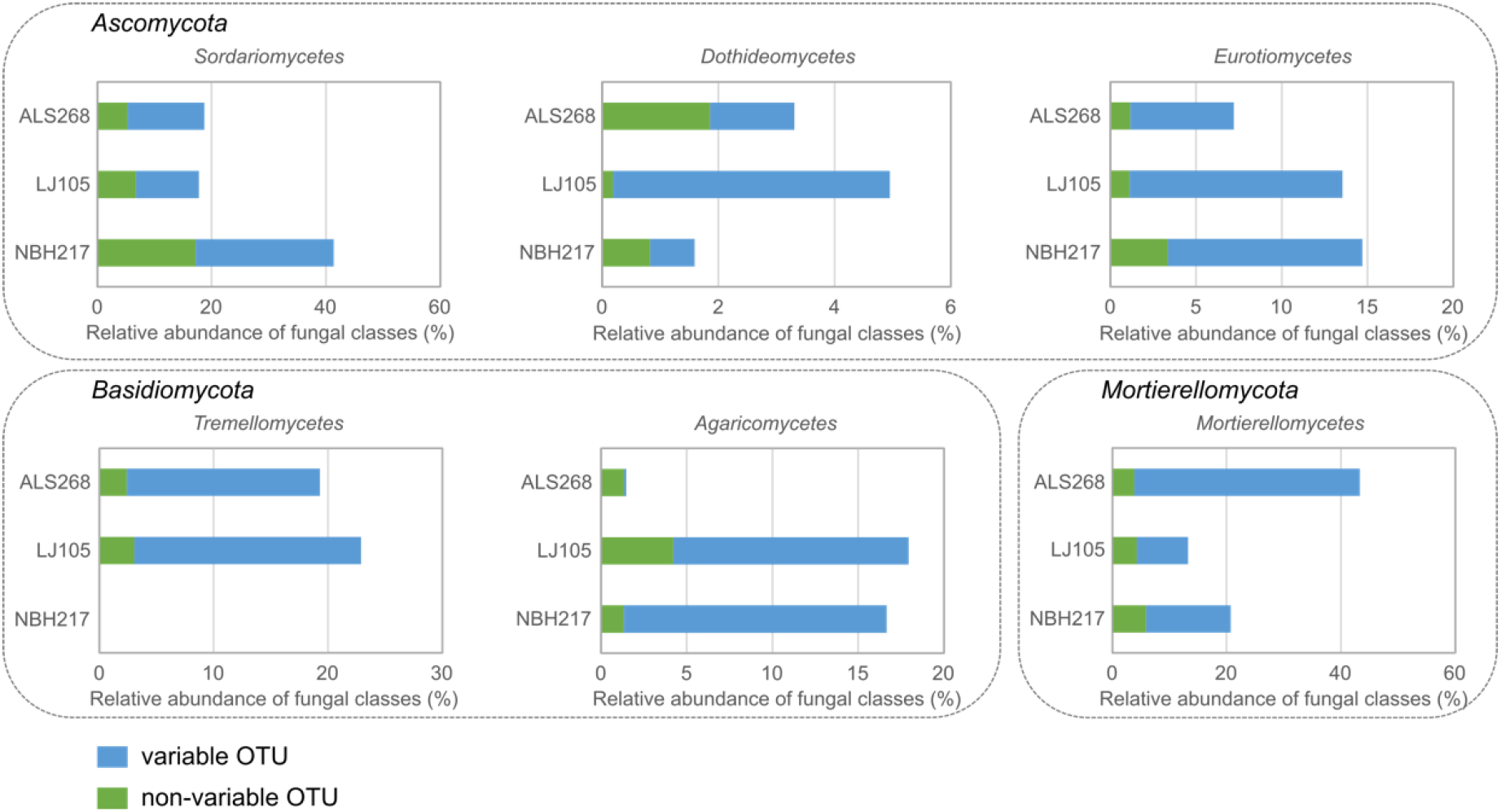
The percentage of variable OTUs (the blue bars) versus non-variable OTUs (the green bars) in different fungal classes for the three technical replicates in all MiSeq runs. ALS=AiLaoShan, LJ=LiJiang, NBH=NaBanHe.

The outcomes showed that run4 and run6 were the main sources of vOTUs, and certain taxa seemed to be associated with the two MiSeq runs (Supplementary Table 3). For instance, the abundance of *Trichoderma barbatum* in all 3 technical replicates was particularly low in runs 4 and 6 but were highly abundant within the other runs. On the contrary, the abundance of *Dactylonectria macrodidyma* in all 3 technical replicates was significantly higher (FDR = 2.1E-05 in ALS, FDR = 4.7E-06 in LJ, FDR = 1.4E-06 in NBH) in runs 4 and 6 than that of the other runs. There were also some taxa with unique distributions across the runs. For example, the abundance of *Sordariales sp.* in ALS268 and *Humicola sp.* in LJ105 were extremely low in run 5 and 8 respectively. In DNBSEQ-G400, only two species from LJ plot sequenced by SE400 mode were characterized as vOTUs, which were also extremely low abundant OTUs, *Penicillium jamesonlandense* (average abundant = 0.33% in all runs of SE400, FDR = 0.005) *and Trichoderma oblongisporum* (average abundant = 0.12% in all runs of SE400, FDR = 0.017) (Supplementary Table 3).

As some of the vOTUs are functional important for soil, this sequencing bias might affect further ecological conclusions drawn from scientific studies. Overall, based on the vOTU results, we found HTS results from the DNBSEQ-G400 platform was more stable than those from the MiSeq platform.

## DISCUSSION

Here, we assessed the amplicon sequencing results of positive controls mock communities and a large number of soil samples DNBSEQ-G400 sequencing platform. Importantly, the cost per sample of DNBSEQ™ is comparable or even lower than that of current platforms (Supplementary Table 1). Since we obtained a high level of reading depth for our testing purposes (~600K reads per sample), the cost for an empirical study, without such high demands in depth, could be further reduced. The DNBSEQ-G400 platform exhibited a high stable performance in our study. As expected, DNBSEQ-G400 could reproduce the relative abundance of designed Zymo mock community, with a similar range of variation as observed for MiSeq based on a previous study (26). These results indicate that the accuracy of the two sequencing platforms are equivalent.

Comprehensive comparisons of ITS2 amplicons from >1000 soil samples revealed highly comparable performance between PE200 and SE400 modes of the DNBSEQ-G400 platform. Specifically, the major fungal classes detected by DNBSEQ-G400 in the PE200 and SE400 modes were similar to that by MiSeq, with differences confined to classes with very low abundance (Fig. 2). However, some important differences between platforms in class abundances were detected. For example, the class Archaeorhizomycetes was highly detected by DNBSEQ PE200 but not by MiSeq. The choice of primers may be the reason for its detection difference, as Archaeorhizomycetes were strongly discriminated against by the ITS4ngsUni primer (27). Further, Archaeorhizomycetes often have highly divergent sequences (27), so the missed detection in the SE400 mode might also suggest the importance of 3’ sequences for certain taxa assignment. Again, the primer differences between the platforms could be the reason for the differences of an unidentified genus of Ascomycota and genus *Mortierella* (Fig. 4b), as the yield of Ascomycota and Mortierellomycota has been reported to be significantly affected by primer pair (10, 28). We initially suspected that variation in sequence read length might also influence the compositional comparisons across platforms. Theoretically, longer reads might improve taxonomic accuracy for fungi with longer ITS2 regions. However, the species that were exclusively detected at MiSeq (PE300) were not the ones with longer ITS2 regions (Supplementary Table 2). Therefore, this possibility is not likely driving the patterns we observed. Moreover, the least similar genera between DNBSEQ-G400 and MiSeq in technical replicates were extremely low in abundance (Fig. 2a), and low abundance genera should be carefully interpreted due to the false positive risk (29).

The evaluation of PCR and run effects on sequencing of technical replicates suggested that DNBSEQ-G400 was more stable than MiSeq. Good consistency among PCR replicates matched with previously reported results, mainly due to the substantial improvements in the processivity and fidelity of DNA polymerase (30). As such, PCR replicates seem to be unnecessary in future projects with possible budget constraints. Instead, we found that sequencing runs have larger effect on the composition of technical replicates than that of PCR, in terms of both MiSeq and DNBSEQ-G400 results (Fig. 4a). However, we only found that there was significant run bias based on fungal composition (Fig. 4a) and/or abundance (Fig. 4b) in the results by MiSeq. Specifically, abundance differences (as determined by vOTUs) were detected in three abundant phyla: Ascomycota, Basidiomycota and Mortierellomycota (Fig. 5). At class level, the shifted proportion of Sordariomycetes and Dothideomycetes in our study were consistent with the previous report on MiSeq run bias against certain fungal taxa (15), suggesting that the biases seems to be group specific. A certain archaeal taxon that was shown to be differentially detected by a Illumina HiSeq run in another run-to-run comparison study further supported this group specific hypothesis (14). Since such bias was observed for several taxa in technical replicates samples, it might also exist in other non-replicated samples throughout the run. While the exact source of run bias remains unclear, cumulative replication errors during signal amplification before sequencing is probably a leading candidate.

The vastly developed sequencing technologies allows deeper discovery of the unknown microorganism in natural complex samples, including shotgun metagenomics and long-read DNA sequencing methods (31–33). However, to reflect the true taxonomic composition of a soil sample still remains a difficult task for scientists, due to the largely undiscovered microorganisms which are lack of references and the complexity of their genomes (34). Compared to metagenomics, metabarcoding could provide more taxonomic information with affordable price, which makes it possible for researchers to expand their surveys on microbial communities to both large spatial and long temporal scales. We presented an alternative platform, which exhibited similar outcomes, but with larger throughput and lower cost, to the currently dominating ones, i.e. Illumina MiSeq. To summarize, the DNBSEQ-G400 platform, which is cheaper on a per base-pair basis and does not exhibit run-to-run bias is recommended for future studies with large sample sizes that requiring high quality control.

## Methods

### Sampling and sequencing experimental design

ZymoBIOMICS Microbial Community DNA Standard (D6305) was bought from ZymoBIOMICS™. Fungus *L. edodes*, *F. velutipes*, *P. eryngii* and *S. cerevisiae* were bought from supermarket as positive controls. Gradient gDNA mixtures were mixed based on DNA content, and then followed the PCR amplification. Meanwhile, each fungus was amplified separately, then mixed according to the content of PCR products to form PCR mixtures.

A total of 1144 topsoil samples collected from forest plots in southwest China were sequenced by both Illumina MiSeq and MGI Tech. DNBSEQ-G400 in this study: 340 soil samples form 20 ha AiLaoShan (ALS) subtropical evergreen broad leaf forest plot, 477 soil samples from 25 ha LiJiang (LJ) subalpine spruce forest plot, and 327 soil samples from 20 ha Nanbanhe (NBH) tropical rainforest plot. The summary information of the 3 plots can be found in Supplementary Table 4. The internal transcriptional spacer 2 (ITS2) amplified individually from each sample were sequenced by both DNBSEQ and MiSeq (Supplementary Fig. 1). All 1144 samples randomly and equally distributed into 3 DNBSEQ runs for both 2×200 paired-end and 400 bases single-end sequencing. The left libraries of total 1144 samples were repetitively sequenced two more times by 400 single end sequencing mode using different DNBSEQ-G400 machine. Similar sequencing experimental design was carried out at Illumina MiSeq platform. Due to the run capacity of MiSeq, 1144 forest soil samples were randomly sequenced by 8 parallel sequencing runs on MiSeq platform. Three randomly chosen soil samples from each forest plot (ALS268, LJ105 and NBH217) were all amplified 3 times separately as PCR replicates. The corresponding 9 sequencing libraries were sequenced repetitively by 3 DNBSEQ runs and 8 MiSeq runs as run replicates (Supplementary Fig. 1).

### DNA extraction

DNA extraction was performed using the PowerSoil®DNA Isolation Kit (Mobio) according to manufacturer’s instructions. DNA concentration was measured by Qubit (Invitrogen).

### Library preparation and sequencing in DNBSEQ-G400

The library preparation for DNBSEQ-G400 was carried out in BGI-Shenzhen. Two-barcode system for metabarcoding on DNBSEQ-G400 platform was designed to allow large input in one sequencing event (Supplementary Fig.2). 15 unique barcodes consisting of 6bp artificial DNA oligos connected with short spacers (0-3 bases random nuclear acid) was linked to 5’ end of ITS1 amplification primers (Supplementary Table 5). Short spacers were used to shift sequences in template DNA to avoid synchronal signals caused by conserved DNA regions in amplicons while sequencing. The first barcode was added to soil DNA through one-step PCR amplification (Supplementary Fig.2). The primers, reaction recipes and conditions for PCR were shown at Supplementary Table 5.

Qualified samples with different barcodes were mixed for DNA nanoball (DNB) based library construction. In this step, second barcodes and sequencing adaptor were added to the mixtures from the first step PCR by ligases. PCRs that labeled by two barcodes were further cyclized, and 20uL of the cyclization fragments were taken to make DNA nanoballs (DNBs). DNBs were sequenced by DNBSEQ-G400 sequencer in BGI-Shenzhen.

### Library preparation and sequencing in MiSeq

Library preparation for Illumina MiSeq was carried out at University of Minnesota Genomic Center according to the following steps. The sequencing libraries were constructed using dual-index (DI) amplification approach (35, 36). The internal transcribed spacer 2 (ITS2) of rDNA was amplified by using two-step polymerase chain reaction (PCR). The primers, reaction recipes and conditions for the two amplifications were shown at Supplementary Table 5. The first PCR products were diluted 100-fold as template DNA for the second amplification. The amplicons were normalized and pooled in equal molar ratios. To ensure the consistency of relative depth, nine PCR reactions of three replicated samples were initially mixed together by equal weight according to the concentration. An equal amount of mixtures was taken out and mixed with the PCR products of other samples when building library for each single run. For each run, all mixtures were purified using Agencourt AMPure XP (Beckman Coulter). The pooled samples were size selected at 427 bp ± 20%, denatured with NaOH, diluted to 8 pM, and then spiked with 15% PhiX. Sequencing was carried out using the entire lane of the Illumina MiSeq platform with a MiSeq 600 cycle v3 kit in the University of Minnesota Genomic Center.

### Processing of ITS2 sequencing data

On average 688.9 million high-quality reads per lane on the DNBSEQ platform were produced using 2×200 paired-end mode, with 88.2% of bases scoring Q30 for read1 and 86.6% bases scoring Q30 for read2. After barcode splitting, an average 558K reads per sample were obtained. For 400 single-end mode in DNBSEQ, an average of 452.5 million high-quality reads per lane were produced, with 79.4% of bases scoring Q30. An average 736K reads per sample were obtained after barcode splitting.

All offline sequencing data was processed using a Python-written Snakemake pipeline. For both paired-end sequencing data (PE200 generated from DNBSEQ-G400 and PE300 from Illumina MiSeq), the forward and reverse reads were initially merged, adaptor and the primer area were removed by cutadapt(37) in the same time. In order to align with UNITE database(38), the 42 bp tails from 3’ end of all sequences were trimmed based on our own test using ITSx(39). Next, low quality reads with maximum Expected Error rate > 1 and minimum length less than 100bp were discarded by Vsearch. Sequences passing these quality filtering steps were separately clustered to OTUs using BURST (https://github.com/knights-lab/BURST) with a similarity of 0.97. We assigned the origin of each read based on its best BURST hit using a ‘winner-takes-all’ strategy which only considers the best match of each sequencing read. Profiles of all samples was therefore concatenated into one OTU table by BIOM (40).

The bioinformatic analysis was slightly modified for single-end sequencing data (SE400) from DNBSEQ-G400. Since SE400 is non-directional sequencing, the direction of sequencing reads was recongnized in the beginning of our analysis. Low quality reads were then removed, and the qualified reads were aligned to database by the same methods used for paired-end sequencing. The assignments of best BURST hits were further produced by ‘winner-takes-all’ strategy, and all profiles were concatenated by BIOM into an OTU table.

All sequencing data from both MiSeq and DNBSEQ-G400 was normalized to 5k sequences per sample by rarefaction, a total of 1000 times. Finally, the samples with less than 5k reads were filtered out, and remaining samples were used in all further analysis.

### Evaluation of the sequencing performance from MiSeq and DNBSEQ-G400 platforms

The similarity of the frequency of occurrences of classes across the sequencing platforms was examined by lme4 package in R (41). Pooled G-test of goodness-of-fit was performed using RVAideMemoire package in R (42). Analysis of similarities (ANOSIM) was performed using Bray distance by vegan package in R (43). Spearman correlation coefficient and the statistics were calculated by vegan and Hmisc functions in R (44). Procrustes analysis was performed by the Procrustes and protest functions from the vegan package in R. The non-metric multidimensional scaling analysis was performed by the metaMDS function from the vegan package in R (43). Nonparametric MANOVA analysis was performed on the abundance of OTUs using Bray-Curtis distance by adonis function from the vegan package in R. Significant differences of the abundance among major OTUs cross sequencing platforms were examined by Kruskal-Wallis test in R. All data was represented by ggplot2 in R (45).

### Identification of vOTUs with significant variation across runs

One-way ANOVA was used to detect fungal species that have significant abundance variation (vOTUs) among runs for all run replicates. To control for a possibly inflated type I error as a result of multiple comparisons, we applied false discovery rate (FDR) testing. Thus, the p value was converted to q value to measure the proportion of FDR using qvalue package in R (46). An OTU with FDR < 0.05 were considered as vOTU.

## Acknowledgements

We thank Prof. Linda Kinkel the help of MiSeq sequencing at University of Minnesota Genomic Center. This research was supported by the National Natural Science Foundation of China (31570380, 31300358, 31100312), the West Light Foundation of the Chinese Academy of Sciences to Yue-Hua Hu, the CAS 135 program (No. 2017XTBG-T01), the Natural Science Foundation of Yunnan Province (2015FB185), the Southeast Asia Biodiversity Research Institute, Chinese Academy of Sciences (2016CASSEABRIQG002), the National Key Basic Research Program of China (2014CB954100), and the Applied Fundamental Research Foundation of Yunnan Province (2014GA003).

## Data Accessibility

The sequencing data for all samples have been deposited to China National GeneBank DataBase (CNGBdb) with the accession number CNP0000069. The authors declare that all other data supporting the findings of this study are available within the manuscript and its supplementary files are available from the corresponding author upon request.

## Author Contributions

X.S., L.X., Y.H. and Z.S. designed the study. Y.H. collected 1144 soil samples from the three forests. X.S., H.M., H.Z., X.L. performed the amplification experiments. J.W., J.L., X.W. and M.G. performed the DNBSEQ-G400 sequencing events. X.S., C.F., Y.J., analyzed the sequencing data. X.S., Y.H. and Z.S. wrote the manuscript.

